# Regulation of input excitability in human and mouse parvalbumin interneurons by Kir potassium channels

**DOI:** 10.1101/2025.04.11.648314

**Authors:** Szabina Furdan, Abdennour Douida, Emőke Bakos, Ádám Tiszlavicz, Gábor Molnár, Gábor Tamás, Daphne Welter, Jonathan Landry, Bálint H. Kovács, Miklós Erdélyi, Balázs Bende, Gábor Hutoczki, Pál Barzó, Vladimir Benes, Viktor Szegedi, Attila Szűcs, Karri Lamsa

**Affiliations:** Hungarian Center of Excellence for Molecular Medicine Research Group for Human neuron physiology and therapy, Szeged, Hungary; Department of Physiology, Anatomy and Neuroscience, University of Szeged, Szeged, Hungary; ELKH-SZTE Research Group for Cortical Microcircuits, Department of Physiology, Anatomy and Neuroscience, University of Szeged, Szeged, Hungary; European Molecular Biology Laboratory, Heidelberg, Germany; Department of Optics and Quantum Electronics, University of Szeged, Szeged, Hungary; Hungarian Centre of Excellence for Molecular Medicine Research Group for Translational Medicine Development, Szeged, Hungary; Department of Neurosurgery, University of Debrecen Clinical Centre, Debrecen Hungary; Department of Neurosurgery, University of Szeged, Szeged, Hungary; Neuronal Cell Biology Research Group, Eötvös Loránd University, Budapest, Hungary

**Author notes:** equal contribution (Szabina Furdan, Abdennour Douida).

## Abstract

Compared to rodents, inhibitory interneurons in the human neocortex exhibit high input excitability because of reduced passive ion leakage across their extracellular membrane. However, the regulation of intrinsic excitability by voltage-gated ion channels activated over a wide range of membrane potentials in human interneurons remains poorly understood. We performed whole-cell patch-clamp microelectrode recordings in mouse and human neocortical slices obtained from surgically resected non-pathological brain tissue finding that Kir channels control the electrical resistance of parvalbumin (Pvalb) neurons in an identical manner in the human and mouse. Molecular analyses revealed predominantly Kir3.1 and Kir3.2 channels in Pvalb neurons in both species. Using whole-cell recordings from synaptically connected neuron pairs and a computational model, we demonstrated that physiological Kir activation inhibits human Pvalb interneurons during postsynaptic potentials evoked by presynaptic neurogliaform cells. The similarity of Kir-mediated inhibition across species suggests that it is an archetypal property of Pvalb neurons.

## Introduction

Rodents are widely used as model organisms in neurobiological research. However, rodent neurons differ from human neurons, limiting the utility of rodent cells for clarifying the molecular and functional characteristics of neurons in the human brain. Human brain tissue, obtained during surgery, serves as a valuable resource for these studies.

Homologous neuronal types (*i.e.*, neurons with similar locations, structures, and developmental origins that express the same marker genes) in the brain exhibit qualitative and quantitative differences in gene and protein expression among mammalian species ^1, 2^. The molecular species characteristics of neuron types are best understood in the neocortex, a component of the cortical mantle involved in complex brain functions ^3, 4, 5^. In addition, studies using *ex vivo* slices prepared from surgically resected human brain tissue have revealed functional differences in the human and rodent neocortical neurons ^6, 7^.

Specifically, human and rodent neocortical neurons differ in the electrical excitability [measured as cell input resistance (Rin)] of the extracellular membrane. Compared with their mouse counterparts, excitatory principal neurons in the human neocortex exhibit reduced responsiveness to electrical currents because of lower Rin ^8, 9, 10, 11^, whereas inhibitory interneurons exhibit increased electrical responsiveness to electrical inputs because of their high plasma membrane resistance ^12, 13, 14, 15, 16^. Input excitability and Rin in human neurons have mainly been studied under resting conditions, *i.e.*, when membrane potential (Vm) is close to −60 mV. However, electrical responsiveness over a wide range of Vm changes, similar to those observed during physiological Vm fluctuations in neurons, is poorly understood in the human neocortex.

Human parvalbumin (Pvalb)-expressing neurons, which comprise the largest inhibitory neuronal population in the cortical mantle ^17, 18, 19^, exhibit higher plasma membrane electrical resistance than their rodent counterparts ^14, 15, 20^. Pvalb neurons induce rapid γ-aminobutyric acid (GABA)-mediated synaptic inhibition of other neurons ^21^ and they are largely conserved across mammalian species. Specifically, Pvalb neurons display similar gene expression of marker molecules (*e.g.*, *PVALB*/Pvalb) and specific ion channels (*e.g.*, *KCNC1*, *KCNC2*) ^22, 23, 24^ and archetypal functional features such as rapid action potentials (APs) and a modestly accommodating AP firing pattern ^21, 25^. However, studies have revealed differences in gene expression, ion channel localization, and electrical function between Pvalb neurons in humans and rodents ^14, 15, 20, 22, 26, 27, 28, 29, 30^.

The voltage-gated ion channel mechanisms controlling intrinsic excitability are well characterized in rodent Pvalb neurons ^31, 32, 33, 34, 35, 36^ but poorly understood in their human counterparts. In rodents, excitability in these cells is strongly regulated by Kir family potassium channels. These channels are active at subthreshold Vm (*i.e.*, below the AP firing threshold), and they effectively inhibit the AP generation ^30, 31, 34^. To close this gap in knowledge, this study investigated the regulation of mouse and human Pvalb neuron excitability by Kir channels in *ex vivo* slices of surgically resected neocortex using whole-cell clamp recordings, patch-sequencing, immunofluorescence labeling, and computational modeling. The inhibitory effect of Kir channels proved identical between human and mouse neurons despite their differences in absolute membrane resistance. Kir-mediated inhibition was progressively strengthened in neurons in both species as the soma Vm became negative relative to the resting potential (−60 mV), causing shunt inhibition of excitatory inputs ^37, 38^. The results revealed robust mRNA expression of Kir3.1 and Kir3.2 channels in human and mouse Pvalb neurons, and confocal and STORM super-resolution fluorescence immunohistochemistry analyses highlighted the expression of the channel proteins in Pvalb neurons. Kir-mediated inhibition was activated in human Pvalb neurons by a physiological mechanism during postsynaptic hyperpolarization caused by AP firing in presynaptic neurogliaform cells (NGFs).

## Results

In this study, human Pvalb neurons (n = 34, see Supplementary Table 1 for cell and patient details) were confirmed to be immunopositive for Pvalb (n = 13) after filling the cells with biocytin or through patch-sequencing to detect *PVALB* mRNA expression (n = 11). If *PVALB* expression was undetectable, then the mRNAs of genes indicative of a neuron type with extremely low Pvalb expression in humans was identified using the Allen Institute Neuron Type Classification System with an algorithm that categorizes neocortex layer 2/3 neuron types by the genes expressed (n = 10; https://knowledge.brain-map.org/mapmycells/process/; Fig. 1a1–2, Supplementary Table 2) ^39^. In parallel, we examined Pvalb neurons in mice using the fluorophore tdTomato in *PVALB*-expressing cells in the brain (n = 10; Fig. 1a3–4).

**Figure 1.**
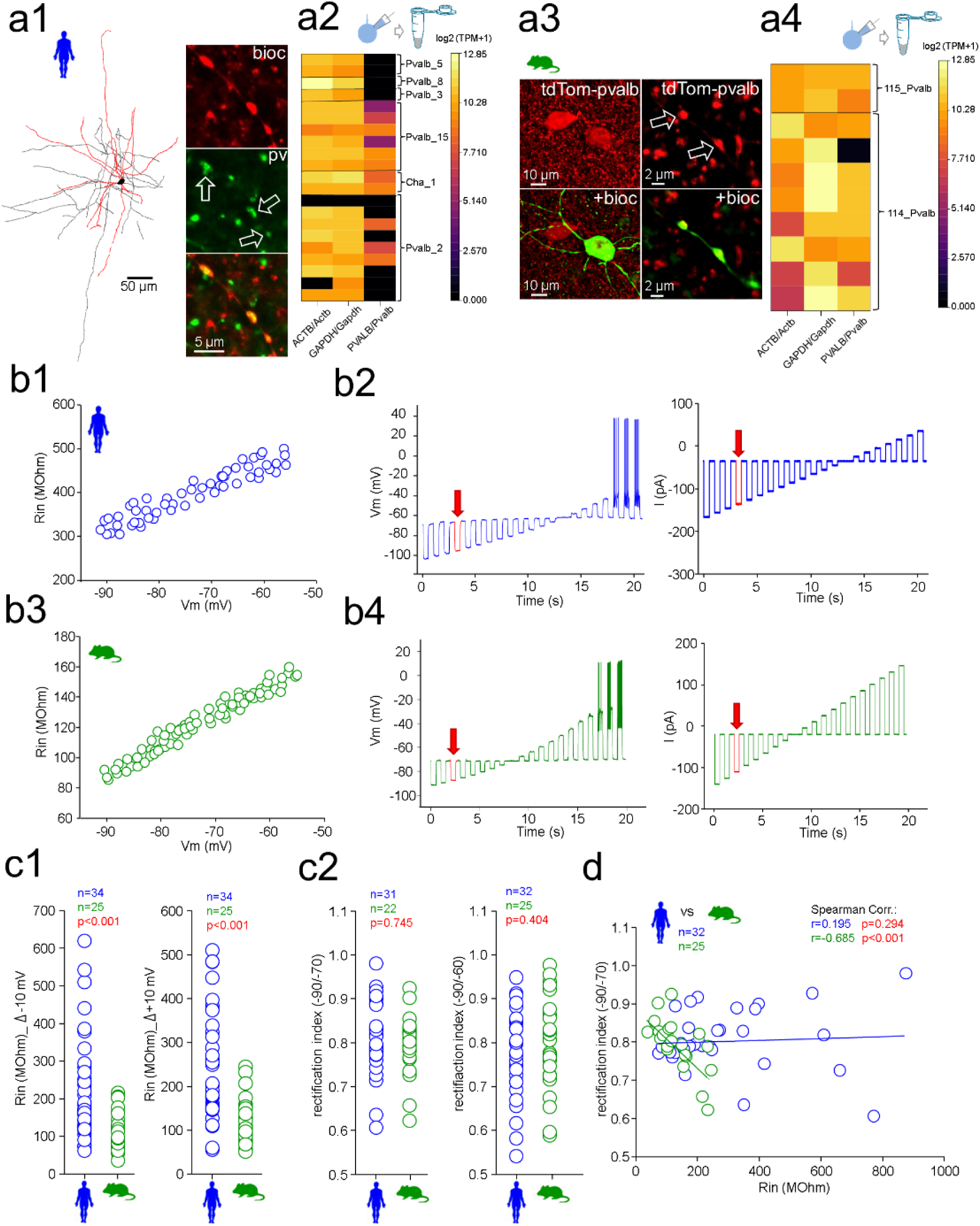
Human and mouse parvalbumin (Pvalb) neurons exhibit identical decreases in input excitability upon membrane potential hyperpolarization despite their differences in input resistance (Rin). **a)** Identification of Pvalb interneurons in neocortex layer 2/3 in humans and mice. **a1)** Some recorded cells were filled with biocytin for *post hoc* visualization of their structure with fluorophore-conjugated streptavidin. The cells were analyzed for the neuronal marker Pvalb by fluorescence immunohistochemistry. *Left:* Visualized human Pvalb neuron after biocytin filling. *Right:* Confocal fluorescence images presenting the Pvalb-immunopositive axon boutons of the same cell. **a2)** Cells were alternatively analyzed for mRNA expression by patch-sequencing. The ordinate presents six Pvalb neuron subtypes identified by their gene expression using the Allen Institute Neocortical Neuron Type Identification System. Some cells showed undetectable *PVALB* mRNA levels. **a3)** Mouse Pvalb neurons were identified using a genetic fluorophore tdTomato (tdTom) in Pvalb neurons. Confocal fluorescence images show the tdTom fluorescence in recorded neuron axon boutons. **a4)** The cells were patch-sequenced and analyzed for mRNA expression. The ordinate presents two mouse Pvalb neuron subtypes as identified by the Allen Institute Neuron Type Identification System. Nearly all cells (9 of 10) showed *PVALB* mRNA. **b)** Incremental amplitude current steps applied to human (**b1–2**) and mouse (**b3–4**) Pvalb neurons reveal a nonlinear current–voltage relationship at subthreshold membrane potential (Vm; *i.e.*, below the action potential generation threshold). The HCN channel blocker ZD7288 (50 μM) is presented in all experiments. **b1)** Rin at different values of Vm in a human Pvalb neuron. **b2)** *Left*: Cellular Vm was hyperpolarized or depolarized at rest (−70 mV) in (*right*) square pulse current steps (500 ms). One step (−100 pA) is highlighted in red and indicated by an arrow to denote the change in Vm. **b3)** Rin at different values of Vm in a mouse Pvalb neuron. **b4)** Vm responses to current pulses in a mouse neuron. A step (−100 pA) is shown in red (arrow) to denote the Vm change in the mouse cell. **c)** Human and mouse Pvalb neurons exhibit similar proportional decreases in Rin upon hyperpolarization even though human cells had a higher absolute Rin. All experiments are in the presence of the HCN channel blocker ZD7288 (50 μM). **c1)** Rin measured by −10 mV (*left*) or +10 mV (right) voltage steps in human and mouse Pvalb neurons. **c2)** Input rectification ratio (r_Rin_) measured at resting potential −60 mV for the ratio of Rin at −90 mV/Rin at −70 mV (*left*) or at −70 mV for the ratio of Rin at −90 mV/Rin at −60 mV (*right*) is similar in human and mouse Pvalb neurons (Mann–Whitney U test). **d)** r_Rin_ is not associated with the Rin in human neurons, but a negative correlation was observed in mouse neurons (Spearman’s correlation test).

### Rin differs between humans and mouse Pvalb neurons but exhibits identical regulation by membrane hyperpolarization

We found that human and mouse Pvalb neurons exhibit a decrease in Rin attributable to Vm hyperpolarization from resting conditions. We applied square-pulse current steps (250–500 ms) at −70 mV in the whole-cell current clamp mode to depolarize and hyperpolarize Vm with voltage steps and systematically measured somatic Rin at steps to −90, −80, and −60 mV. As illustrated in Fig. 1b1–4, Rin at all Vm values was higher in human neurons than in mouse cells (−90 mV: 238.8 ± 171.8 MΩ vs. 103.1 ± 40.8 MΩ; −80 mV: 256.2 ± 183.4 MΩ vs. 115.5 ± 44.5 MΩ; −60 mV: 291.9 ± 206.1 MΩ vs. 135.2 ± 53.3 MΩ; Fig. 1c1).

However, human and mouse Pvalb neurons exhibited similar proportional decreases in Rin in response to hyperpolarizing Vm. Rin in human Pvalb neurons was 18.2% lower at −90 mV than at −60 mV, whereas a difference of 23.8% was observed in mice. The resulting input rectification ratio (r_Rin_; Rin at −90 mV/Rin at −60 mV) was 0.77 ± 0.09 in humans, versus 0.81 ± 0.08 in mice (p = 0.404; Fig. 1c2). In addition, we measured Rin in the same cells by applying only negative current steps (up to −90 mV) at −60 mV, recording a 19.3% decrease in Rin in humans (r_Rin_ [−90/−70 mV] = 0.81 ± 0.08) and a 22.6% decrease in mice (r_Rin_ [−90/−70 mV] = 0.80 ± 0.07; Fig. 1c2). The experiments were performed in the presence of an HCN current blocker (ZD7288, 50 μM) ^15^.

r_Rin_ was not significantly associated with the initial Rin in human cells (r = 0.195, p = 0.294), but higher Rin was associated with stronger r_Rin_ in mouse cells (r = −0.685, p < 0.001; Fig. 1d).

### The voltage-dependent regulation of somatic Rin is mediated by Kir-type potassium channels in human and mouse Pvalb neurons

Next, we investigated the mechanisms underlying the decrease in cell Rin associated with Vm hyperpolarization in Pvalb neurons. Ba^2+^ (100 μM) applied via extracellular solution reduced the voltage dependence of Rin in human (Fig. 2a1–2) as well as in mouse (Fig. 2b1–2). The progressive decrease in Rin at incrementally hyperpolarized Vm values was reduced in the presence of Ba^2+^ in both species, and Rin displayed a more linear relation at Vm values of −90 to −60 mV. r_Rin_ in human cells increased from 0.78 ± 0.02 (baseline) to 0.88 ± 0.05 in the presence of Ba^2+^(p = 0.022). Similarly, r_Rin_ in mouse Pvalb cells increased from 0.81 ± 0.02 to 0.90 ± 0.06 after the addition of Ba^2+^ (p = 0.009) (Fig. 2c1–2). As summarized in Fig. 2, the effect of Ba^2+^ on r_Rin_ was similar between human and mouse Pvalb neurons (p = 0.673).

**Figure 2.**
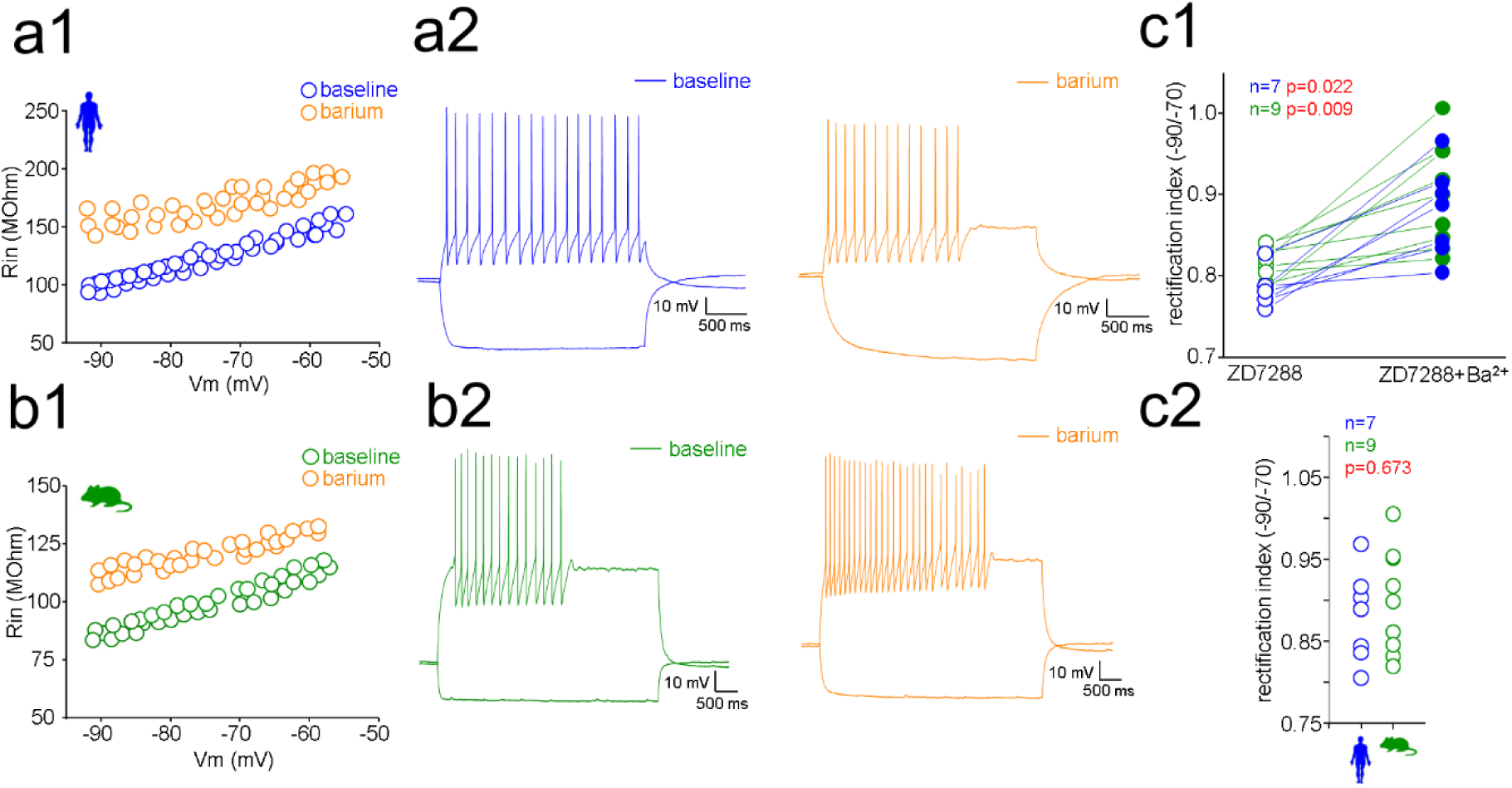
Parvalbumin (Pvalb) neuron input resistance (Rin) decrease induced by hyperpolarization is reduced by Ba^2+^ in humans and mice. Extracellular Ba^2+^ (100 μM) suppresses the decrease in Rin induced by hyperpolarization (HCN channel blocker ZD7288, 50 μM is presented in all experiments). **a1)** Rin in a human Pvalb neuron measured at different membrane potentials (Vm) under control baseline conditions (blue) and in the presence of extracellular Ba^2+^ (orange, 10 min). **a2)** Vm steps under control baseline conditions (green) and in the presence of Ba^2+^ (orange). **b1)** Rin in a mouse Pvalb neuron at different values of Vm under control baseline conditions and in the presence of Ba^2+^. **b2)** Traces presenting Vm steps under both conditions. **c1)** Plot presenting Input rectification ratio (r_Rin_) in human (blue) and mouse (green) Pvalb neurons under control baseline conditions and following exposure to Ba^2+^. r_Rin_ was similarly increased by Ba^2+^ in human and mouse (Wilcoxon’s test). **c2)** Plot shows r_Rin_ in the presence of Ba^2+^ in the human and mouse (Mann–Whitney U test).

The progressive reduction of Rin at hyperpolarized Vm values and its blockade by micromolar Ba^2+^ levels indicate that the decrease in Rin is mediated by the activation of inwardly rectifying potassium currents (*i.e.*, Kir channels) during hyperpolarization ^31, 38, 40^.

### Human and mouse Pvalb neurons exhibit high mRNA expression of Kir3.1 and Kir3.2 channels

We used patch-sequencing to examine the mRNA expression [transcripts per million (TPM)] of genes encoding Kir potassium channel types in human and mouse Pvalb neurons ^16^. Among 16 different *KCNJ* genes (*KCNJ1*–*16*) encoding Kir potassium channels, patch-sequencing in human cells revealed the strongest mRNA expression for *KCNJ3* and *KCNJ6* encoding Kir3.1 and Kir3.2 channels, respectively (Fig. 3a). Kir3.1 and Kir 3.2 mRNA expression was also detected in mouse *Pvalb*-expressing neurons, albeit at lower levels for *KCNJ3* (p = 0.012) but at similar levels for *KCNJ6* (p = 0.07) relative to the findings in human neurons. In addition, human and mouse Pvalb neurons displayed differences in the expression of *KCNJ4* (encoding Kir2.3; abundant in humans but absent in mice, p = 0.018) and *KCNJ9* (encoding Kir3.3; abundant in mice but weakly expressed in humans, p = 0.03).

**Figure 3.**
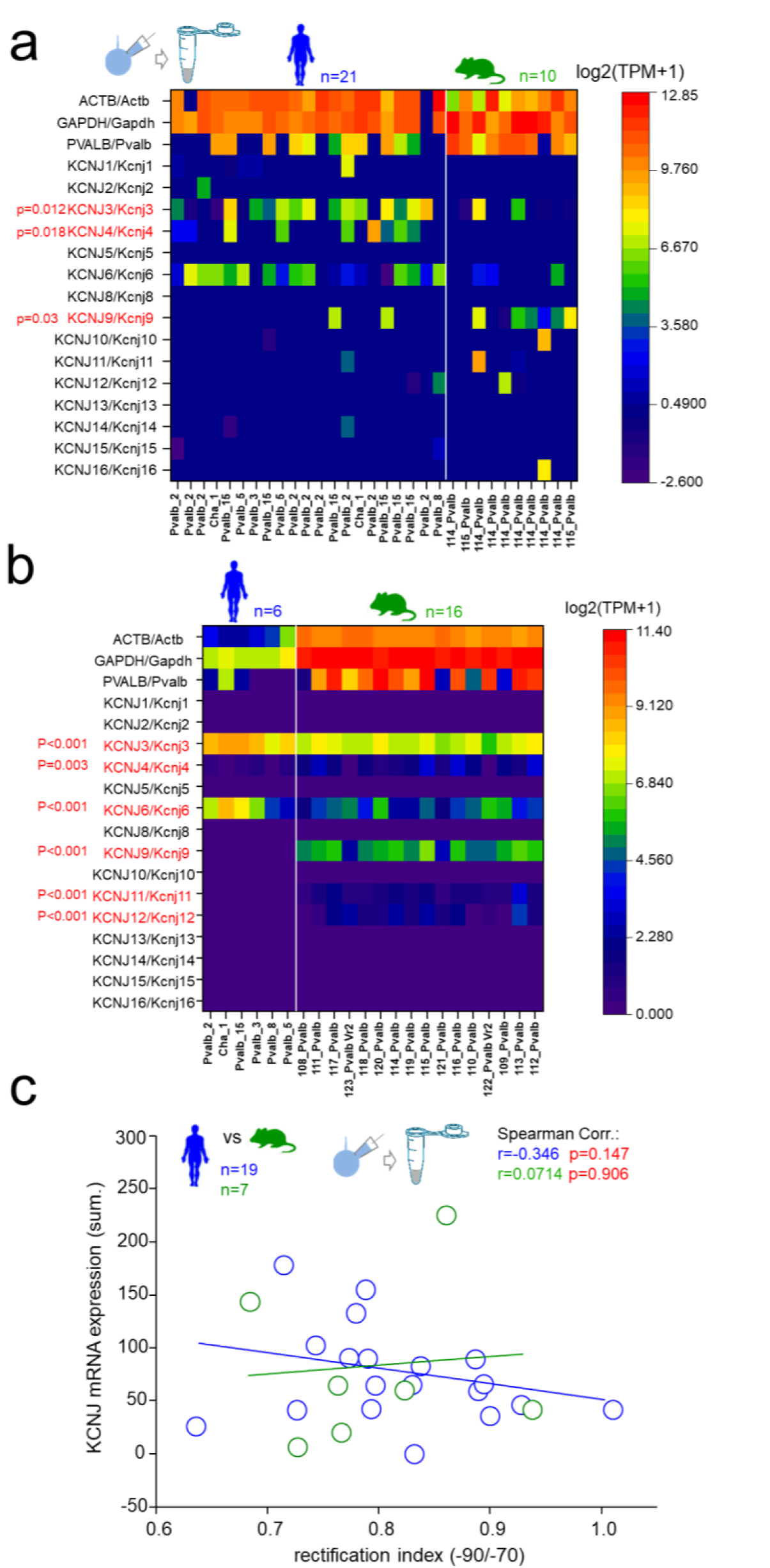
Human and mouse parvalbumin (Pvalb) neurons express high levels of Kir3.1 and Kir3.2 mRNA.. **a)** Analysis of mRNA in patch-sequenced human and mouse Pvalb neurons. The heatmap presents the mRNA levels (transcripts per million [TPM]) in individual human (n = 21) and mouse (n = 10) neurons for the genes *KCNJ1–16* encoding Kir family channels, as well as the housekeeping genes *GAPDH* and *ACTB*. The abscissa presents Pvalb neurons with their subtype identity defined by their gene expression pattern in the Allen Institute Neocortical Neuron Type Identification System (Pvalb neuron type identity codes shown in the abscissa). Human and mouse cells exhibit mRNA of *KCNJ3* (Kir3.1) and *KCNJ6* (Kir3.2), and also *KCNJ9* (Kir3.3) which is higher in mouse. In addition, the human cells express *KCNJ4* (Kir.2.3). p-values for the difference between human and mouse are presented on the left (MANOVA with Bonferroni’s *post hoc* test) with significance difference denoted by red text. **b)** The heatmap presents the Allen Institute data for the mean *KCNJ* gene mRNA level (TPM) in the different Pvalb neuron subtypes in humans and mice. The mRNA levels are high in human and mouse cell types for *KCNJ3* and *KCNJ6*. Mouse cells have higher expression of *KCNJ4*, *KCNJ9, KCNJ11*, and *KCNJ12* (MANOVA with Bonferroni’s *post hoc* test). **c)** The plot presents the lack of an association between the Kir channel gene mRNA level (TPM, summed for all *KNCJ* types) and input rectification ratio (r_Rin_). r_Rin_ = 1 indicates the lack of a Kir channel effect (Spearman’s correlation test).

In addition, we examined the average transcriptomic profiles of Kir channel genes in the human middle temporal gyrus as determined by the Allen Brain Institute ^5, 41^. The analysis revealed high expression of Kir3.1 and Kir 3.2 in both species (Fig. 3b), in line with the results of patch-sequencing. Both genes exhibited higher expression in human neurons than in mouse neurons (both p < 0.001). In addition, *KCNJ9* expression was robust in mice but absent in humans (p < 0.001), as also observed via patch-sequencing. Conversely, *KCNJ4* was detected in mice but not in humans (p < 0.001), contradicting the results of patch-sequencing. The data also demonstrated a significant difference in TPM for *KCNJ11–12* (both higher in mice, p < 0.001), which was not revealed via patch-sequencing.

The total mRNA expression (TPM for all Kir-type mRNAs summed in a cell) in neurons was not associated with the strength of the hyperpolarization-associated decrease in Rin in humans (r = 0.346, p = 0.147) or mice (r = 0.0714, p = 0.906; Fig. 3c). The total level of *KCNJ* mRNAs (summed TPM) was not associated with the absolute Rin measured at −70 mV (human: r = −0.384 p = 0.105; mouse: r = −0.178, p = 0.713).

### Kir3.1 and Kir3.2 channel proteins are present in human and mouse Pvalb neurons

We conducted immunofluorescence analysis of the two most abundantly expressed Kir channel types, namely Kir3.1 and Kir3.2, in human and mouse neocortical sections. Double immunofluorescence staining for Pvalb and either Kir3.1 or Kir3.2 and their analysis in parallel by confocal and STORM super-resolution fluorescence microscopy ^15^ uncovered the localization of Kir3.1 and Kir3.2 in the cells.

Subsequently, quantitative analysis of Kir3.1 and Kir3.2 based on Pvalb immunoreactivity (or tdTomato fluorescence in mice) was conducted utilizing confocal images of human and mouse neurons (Fig. 4a–d). Each line was used to measure the Pvalb (or tdTomato in mice) fluorophore emission in parallel with the Kir3.1 or Kir3.2 immunofluorescence intensity over a distance of 3 μm from the extracellular space, and across the extracellular membrane and 5 μm towards the center of the cell soma (see Fig. 4a1-2). Human neurons exhibited Pvalb immunofluorescence within the soma, whereas mouse cells demonstrated intense somatic emission of tdTomato, a transgenic fluorophore driven by the *PVALB* promoter. In the line analysis, the plasma membrane area manifested as a narrow zone in which the intracellular fluorescence signal rapidly appeared in human (Pvalb) (Fig. 4a1–2) and in mouse (tdTomato) (Fig. 4b1–2) ^15^. The immunofluorescence intensities of the labeled Kir3.1 and Kir3.2 channels were subsequently measured and quantified along the line with the Pvalb (or tdTomato) signal.

**Figure 4.**
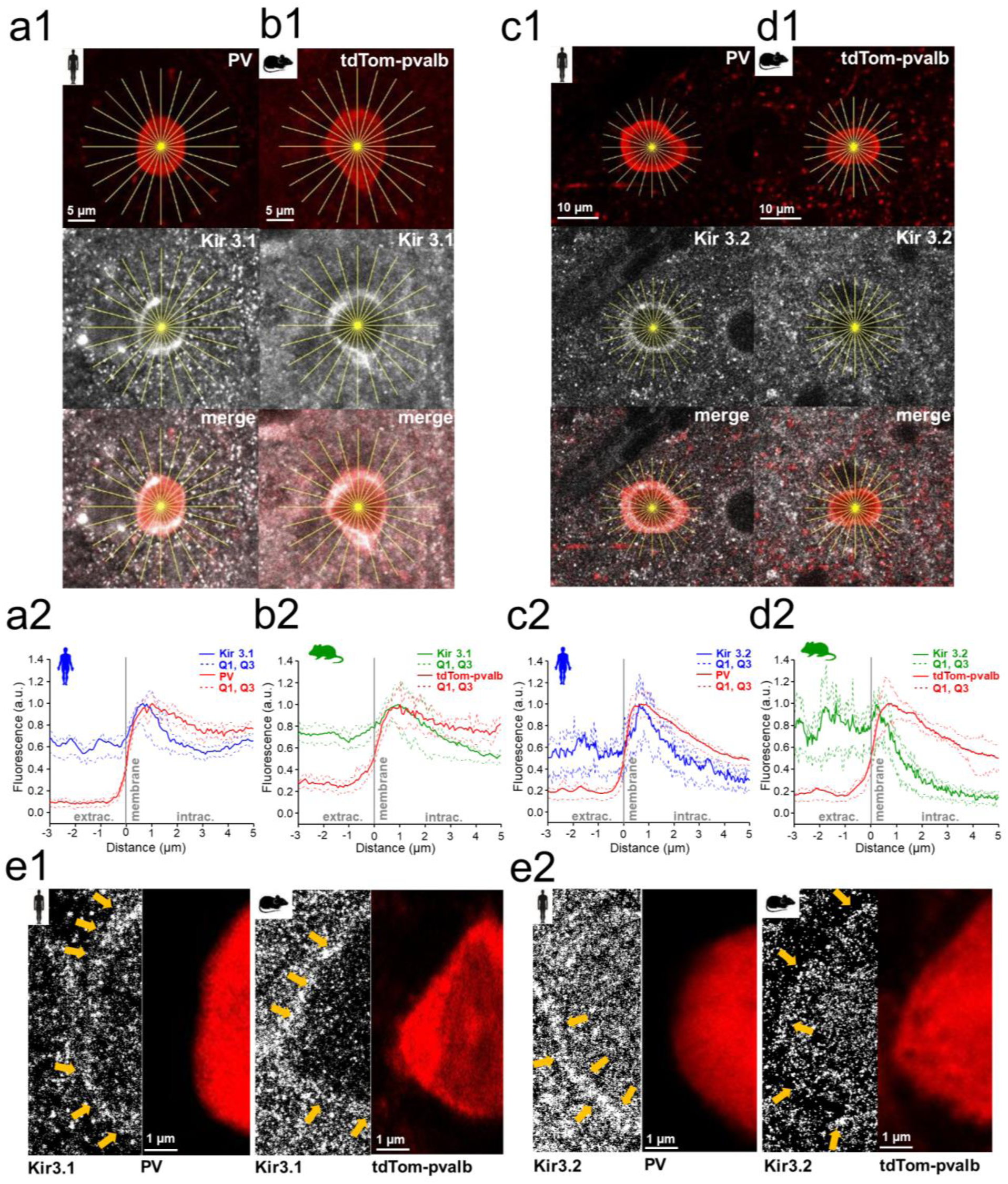
Kir3.1 and Kir3.2 channel proteins are present in the soma of human and mouse Pvalb neurons. Immunofluorescence analysis with confocal microscopy for Kir3.1 or Kir3.2 channel proteins in parallel with the reaction for parvalbumin (Pvalb; or Pvalb-linked tdTomato in mice) illustrates the presence of Kir proteins in the cytoplasm and putative plasma membrane of the soma in human and mouse cells. Panels **a–d** present data for a similar experiment in human (**a1–a2**) and mouse neurons (**b1–b2**) with Kir3.1 and Pvalb (or tdTomato) and in human (**c1– c2**) and mouse neurons (**d1–d2**) with Kir3.2. **a1)** Confocal microscopic image of immunofluorescence reactions in Pvalb neurons (top), in layer 2/3 of the human neocortex for Kir3.1 (middle), and their merged images (bottom). The immunofluorescence intensity for Pvalb (PV) and Kir3.1 was measured in a radial pattern of lines (presented in yellow in the images) that diverged from the extracellular space and projected into the center of the cell soma. Immunofluorescence intensity was measured along 24 radial lines. **a2)** Immunofluorescence intensity (ordinate, arbitrary units) for both Pvalb (PV, red line) and Kir3.1 (blue line) measured along lines from the extracellular space (negative abscissa values) to inside the cell soma (positive values) crossing the extracellular membrane (0 point). The extracellular membrane (vertical dashed line) in the lines is determined by the appearance of the Pvalb immunofluorescence signal. Solid and dashed lines denote the median and quartiles (Q1 and Q3) of the intensity values in 24 pooled lines, respectively. The blue line denotes the peak intensity at the extracellular membrane region. **b1)** Confocal fluorescence image of tdTomato (top) in Pvalb neurons in layer 2/3 of the mouse neocortex, image of immunoreactivity for Kir3.1 (middle), and the merged image (bottom). **b2)** Pooled data from 24-line analyses of tdTomato (red) and Kir3.1 fluorescence (green) in mouse cells (median and interquartile range). The Kir3.1 intensity peak is located at the extracellular membrane. **c1)** Pvalb and Kir3.2 immunoreaction line analysis in human neurons. **c2)** Pooled data from the 24-line analysis of the fluorescence intensity. The Kir3.2 immunofluorescence peak in human cells is located at the extracellular membrane. **d1)** TdTomato and Kir3.2 confocal images in mouse cells. **d2)** Median and quartiles of the 24-line analysis. Kir3.2 immunofluorescence peak is at the extracellular membrane in mouse Pvalb neurons. **e)** Black and white micrographs present dSTORM super-resolution images of the immunofluorescence reaction (at Cy5) for Kir 3.1 (**e1**) and Ki3.2 (**e2**) with confocal microscopic images of Pvalb (Cy3, human) and tdTomato (mouse). Arrow lines in the Storm images indicate Kir3.1 or Kir3.2 antibody locations on the extracellular membrane (compared with cytoplasmic Pvalb immunofluorescence).

Membrane localization was observed in the line analysis of human (Fig. 4a1–2) and mouse cells (Fig. 4b1–2) as the Kir3.1 fluorescence intensity peak at the site at which the immunofluorescence for Pvalb (human) or tdTomato (mouse) rapidly appeared. Similar measurements of Kir3.2 immunofluorescence demonstrated its presence at the extracellular membrane of the soma in both species (Fig. 4c1–2 and 4d1–2). The Kir3.2 immunofluorescence peak in line analyses was also observed at the site at which the somatic fluorescent markers (Pvalb in human; tdTomato in mouse) rapidly appeared (Fig. 4c2 and 4d2, respectively).

Figure 4 presents the STORM immunofluorescence images for Kir3.1 and Kir3.2 channel proteins in selected Pvalb neurons in human (Fig. 4e1) and mouse (Fig. 4e2). The images highlight the localization of individual fluorophore molecule signals at the extracellular membrane.

Computational single-cell model illustrates that stronger somatic transmembrane Kir conductance (G_Kir_) is required for excitability regulation in mouse neurons than in human neurons

We used a computational single-cell model to mimic the Vm dynamics of Pvalb neurons recorded in whole-cell. For simulations with the computational model, we selected a Pvalb neuron (Fig. 5a1–2) with intrinsic electrical properties observed both in human as well as mouse neurons (neuron H4, Supplementary Table 1) when input currents were applied to the soma. The model Pvalb neuron (Fig. 5b1—2) had a soma with passive voltage-independent leak membrane conductance (G_leak_), HCN-type currents, Kv7- and Kv1-type potassium currents in the soma and dendrites, and AP generating Na currents in the axon ^42^. The Rin and capacitance of the model cell mimicked the real neuron, and the Vm changes produced by the input currents in the model cell (from −300 pA to +200 pA square pulse in 250-ms steps) replicated the real Pvalb neuron (Fig. 5c1–2).

**Figure 5.**
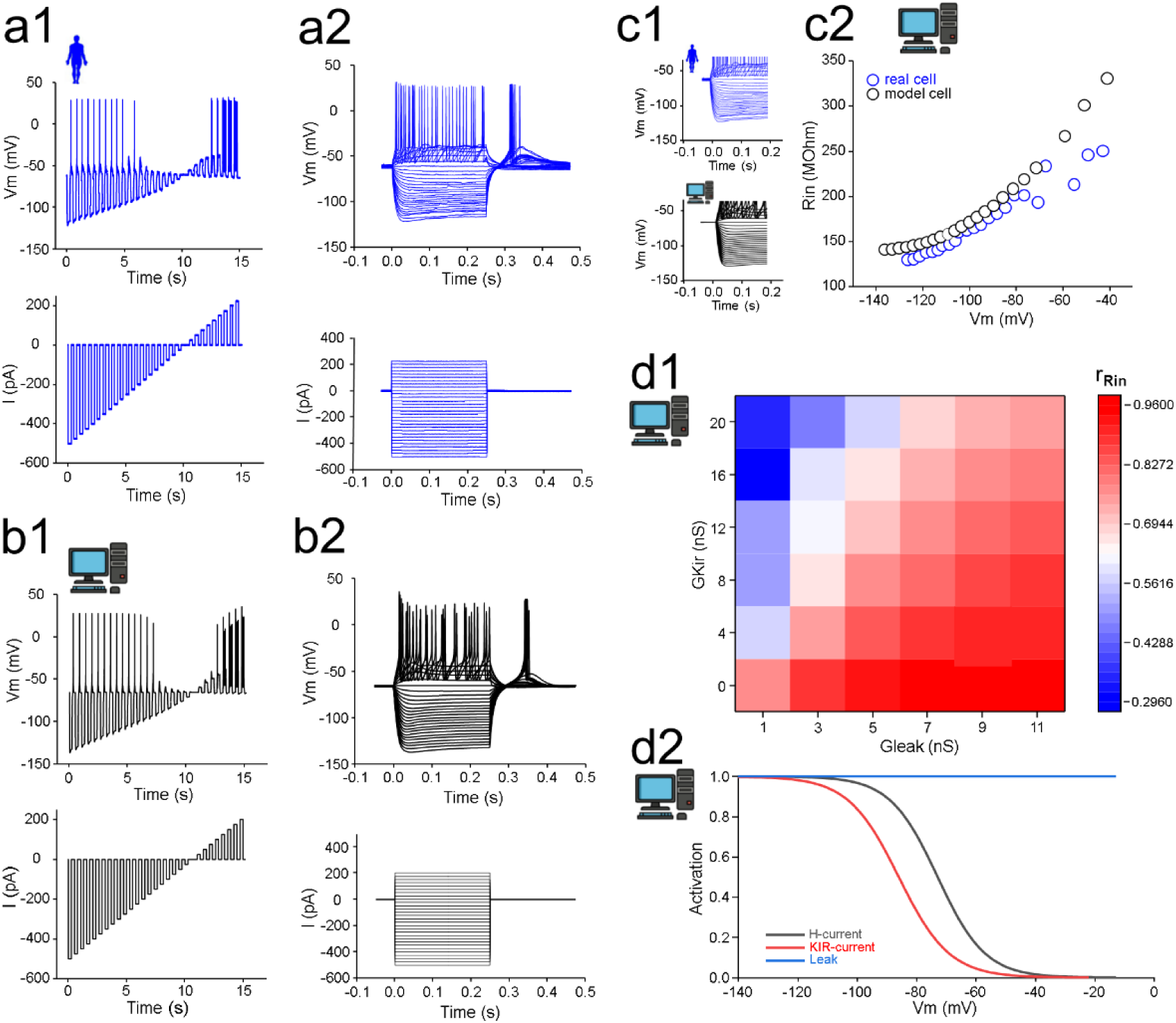
Computational model simulation mimicking the regulation of input excitability by Kir channel activity in parvalbumin neurons with differences in passive leak membrane conductance (G_leak_). **a)** Recording from a Pvalb cell in the human neocortex. **a1)** The I-V step protocol show Vm changes by different current (I) -steps. **a2)** Superimposed Vm step traces (top) and the current steps (bottom). **b)** Single-cell computational model with passive and voltage-activated electrical conductance in the soma membrane mimics the real cell Vm responses through I steps. **b1)** The I-V step protocol shows Vm changes by different I-steps. **b2)** Superimposed Vm step traces (top) by the current steps (bottom). **c)** The Pvalb model cell (black symbols) mimics Rin at different values of Vm measured in the real neuron (blue). **c1)** Superimposed Vm traces in the real Pvalb neuron (blue) and the model cell (black). **c2)** The plot presents Rin at different values of Vm in the real (blue) and model cells (black). **d)** Soma input rectification ratio (r_Rin_; measured from I-V steps as Rin at −90 mV/Rin at −70 mV) in the model cell simulation with variable G_leak_ (abscissa) and Kir conductance (G_Kir;_ ordinate). The cell has a constant small gHCN (0.4 nS at −70 and 0.9 nS at −90 mV) as occurs in real cells ^15^. **d1)** The heat map presents r_Rin_ in the model cell in different configurations when G_leak_ in the soma varied from 1 to 11 nS and G_Kir_ varied from 0 to 20 nS. **d2)** Because G_Kir_ depends on Vm, the simulated cell has an active G_Kir_ (steady-state red line) of 10% of the maximal activation value G_Kir_ (shown in ordinate) at −70 mV and 60% at −90 mV. G_leak_ (blue line) is independent of Vm. The model cell has gHCN (black line) of 1 nS (maximum) with the steady-state voltage dependence between 04 to 0.9 nS presented in the plot ^15^.

To determine the actual Kir current conductance required for r_Rin_ observed in whole-cell recordings from real neurons over a wide range of resting Rin, we simulated the model cell with variable G_leak_ in the soma between 1 and 11 nS. In addition, we simulated the cell with variable Kir channel conductance (ranging from 0 to 20 nS maximum G_Kir_ localized in the soma of the model cell) (Fig. 5d1). Because G_kir_ is voltage dependent, this yielded to a steady-state activation level of approximately 10% of the maximum G_Kir_ at −70 mV, and 60% at −90 mV (Fig. 5d2).

The range of resting Rin values in the simulation covered values measured in real human and mouse Pvalb neurons during whole-cell recordings, and the range of G_kir_ was adjusted to ensure that r_Rin_ corresponded to the findings in actual human and mouse Pvalb neurons. The heat map in Fig. 5d1 presents the voltage dependence of the intrinsic excitability of the model Pvalb neuron under these conditions, and the color coding indicates r_Rin_ (Rin at −90 mV/Rin at −70 mV). The simulation shows that r_Rin_ in neurons with lower G_leak_ requires less Kir channel conductance in somata to generate similar r_Rin_ as that generated in cells with high G_leak_. Fig. 5d2 presents the steady-state activation profiles of Kir current conductance in the simulations, G_leak_, as well as the HCN current conductance ^15^.

### Physiological regulation of input excitability by Kir channel activation in human Pvalb neurons

The recordings in Fig. 6a1–3 present three separate whole-cell experiments from monosynaptically connected pairs of neurons, identified as putative presynaptic NGFs and postsynaptic fast-spiking interneurons, in the human neocortical slices. The experiments illustrated that the APs (two APs delivered 50 ms apart) in presynaptic NGFs (soma in layer 1) evoked slow-kinetic and long-lasting Vm hyperpolarization in postsynaptic fast-spiking interneurons in the layer 2/3 (Fig. 6a1–3). A slow and sustained hyperpolarizing inhibitory postsynaptic potential (IPSP), mediated by GABA-B receptor activation and Kir channels, is a hallmark of NGF firing in the rodent and human layer 2/3 neurons, including fast-spiking interneurons ^12, 43, 44, 45, 46^. In response to Vm hyperpolarization, the postsynaptic fast-spiking neurons (Fig. 6a1–3) exhibited a Kir-type channel-mediated decrease in Rin similar to that observed for r_Rin_ in Figures 1–2.

**Figure 6.**
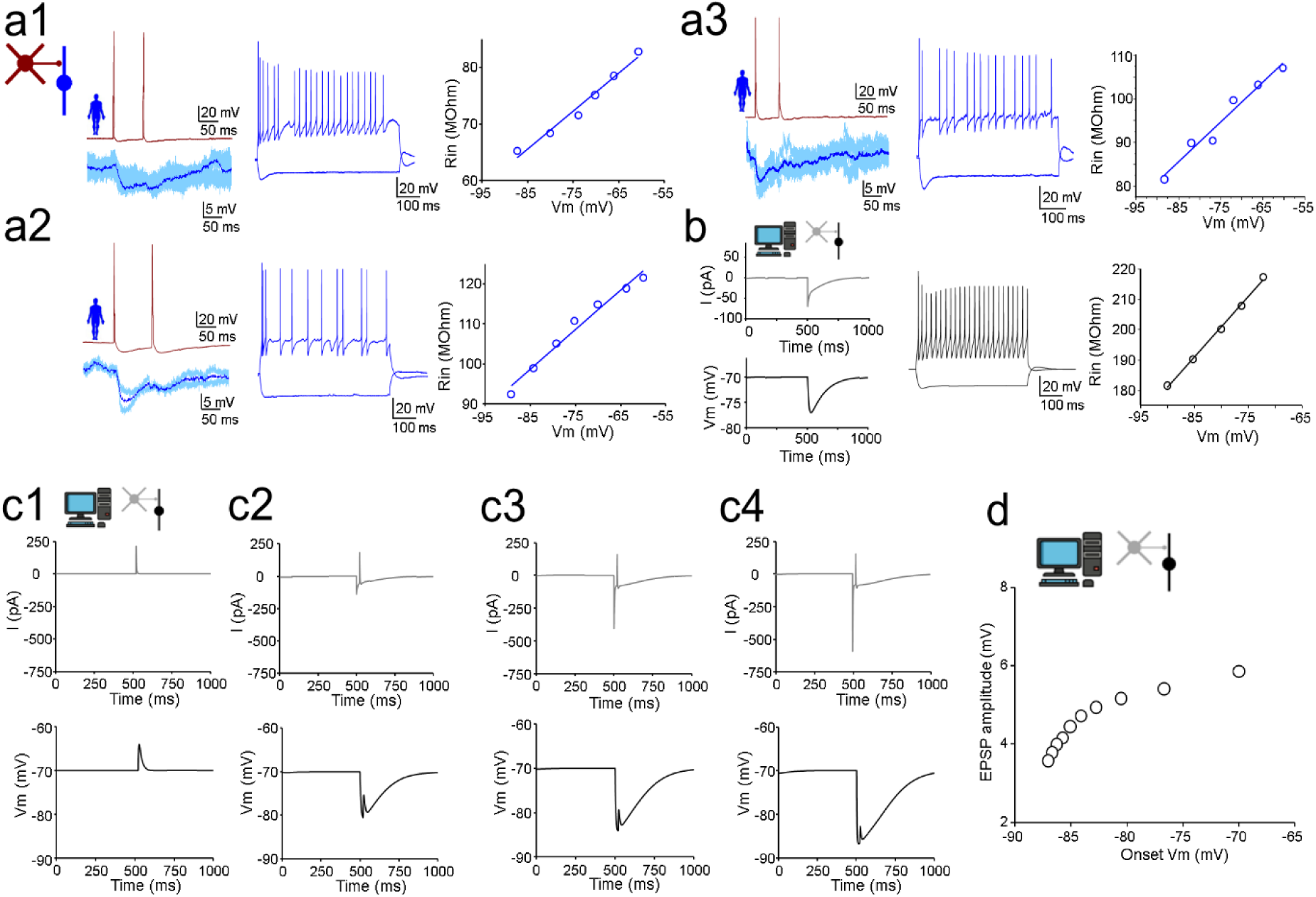
Kir current activation decreases input excitability in human fast-spiking interneurons during synaptic inhibition from neurogliaform cells (NGFs). **a)** Three whole-cell recording experiments **(a1-a3)** of monosynaptically connected pairs with presynaptic putative NGFs and postsynaptic fast-spiking interneurons in the human neocortex. Action potentials (APs) in presynaptic NGFs generate a slow and sustained Kir-mediated inhibitory postsynaptic potential (IPSP) in the postsynaptic fast-spiking interneuron soma. **a1– a3)** *Left*: APs in presynaptic (brown) NGFs (evoked by two 5-ms depolarizing pulses separated by 50 ms) generate monosynaptic IPSP characterized by a long decay (blue traces, bold trace shows the mean). The inset in (a1) presents the schematic of the experiment. *Middle*: AP spiking pattern of postsynaptic fast-spiking interneurons during a depolarizing I-step. *Right*: Input resistance (Rin) is reduced by hyperpolarization in the postsynaptic fast-spiking cells. **b)** Simulation of the Pvalb neuron model cell mimics the IPSP generated by presynaptic NGFs. The inset presents the experimental scheme. *Left*: Traces from the simulation of the Pvalb model cell with a slowly decaying hyperpolarizing Vm change, mimicking the monosynaptic IPSP. *Middle*: The firing pattern of the model cell during a depolarizing I-step. *Right:* The Rin decrease attributable to Vm hyperpolarization. **c)** Pvalb model cell presenting the Kir-mediated inhibition of excitatory postsynaptic potentials (EPSPs) by IPSPs (mediated by Kir activity) from an NGF. **c1–c4)** Simulation of fixed-strength EPSCs (top) in soma generating EPSPs (bottom) in the model cell. In addition, a Kir-mediated IPSP with increasing IPSC amplitude (top) is generated simultaneously with the EPSP (bottom). The IPSP increases from its absence (**c1**) to reach a negative peak Vm at −80 (**c2**), −85 (**c3**), and −87 mV (**c4**; resting potential, −70 mV). (D) Plot shows that the EPSP amplitude (from the onset to the peak) is progressively diminished at more hyperpolarizing values of Vm by the Kir-mediated IPSPs.

We mimicked a hyperpolarizing IPSP in the Pvalb model neuron (Fig. 6b) by Kir channel activation, as estimated from the Rin voltage dependence in the model cell. Finally, we simulated excitatory postsynaptic potentials (EPSPs) in the soma of the model Pvalb neuron (Fig. 6c1–4). EPSPs were generated by simulating excitatory postsynaptic currents (EPSCs) with a peak strength of 3 nS in the soma during the time course of the IPSP generated by Kir channel activation (Fig. 6c1–4). The results showed that during IPSP, the EPSP amplitude is reduced because of shunt inhibition by decreased Rin through Kir channels (Fig. 6d) ^37^.

## Discussion

The regulation of input excitability across a wide physiological Vm range is poorly understood in many neuron types in the human brain. We found that input excitability in Pvalb-type inhibitory neurons in the human neocortex is regulated by Kir family potassium channels. The Kir channel-mediated inhibitory effect (measured as the relative change in Rin) was identical in human and mouse Pvalb neurons, although Rin was higher in absolute terms in human cells. Kir channel-mediated excitability inhibition was similarly activated in human and mouse neurons upon incremental hyperpolarization from resting Vm, although both species exhibited considerable cell-to-cell variability in the strength of the effect.

The similarity of Kir-mediated inhibition between these species, despite their differences in electrical Rin, shows that this regulatory mechanism in the soma is conserved in Pvalb neurons in the mammalian neocortex. The human neurons achieve excitability inhibition in the soma at a lower Kir channel conductance density in the plasma membrane than their rodent counterparts ^47^. This is neuroeconomically advantageous for human neurons, because it reduces the work of maintaining ion gradients across the extracellular membrane in these highly active and metabolically demanding cells ^48, 49^. Kir-mediated inhibition in the soma is predominantly mediated by Kir3.1 and Kir3.2 channels in both species. Physiological Kir activation inhibits Pvalb interneurons during inhibitory postsynaptic potentials (IPSPs) evoked by a presynaptic neurogliaform cell.

The results demonstrate a conserved evolutionary feature of a major ion channel mechanism in mammalian Pvalb neurons, while certain other channels are differentially present in these cells in humans and rodents ^15, 16^. The current study focused on Kir channels and was restricted by the limited availability of resected brain tissue for electrophysiological and molecular studies. Further studies are needed to gain a better understanding of the ion channels in analogous neuron types in humans and laboratory animals.

### Kir-mediated excitability control is part of the archetypal Pvalb interneuron

Certain functional properties of Pvalb neurons are conserved in different mammalian species ^25^. An evolutionarily conserved Kir channel-mediated excitability regulation is not self-evident, because for example, other ion channel mechanisms, such as HCN or Kv1-type potassium channels, contribute differently to excitability control in human and rodent Pvalb neurons ^15,16^.

One explanation for the conserved effect of Kir channels is their strong inhibitory action efficiently combining voltage inhibition and shunt inhibition in the soma. This rapidly suppresses input–output transformation in Pvalb neurons^31, 37, 49, 50^ and prevents the generation of excessive AP discharges in these cells, which are vulnerable to damage from epileptiform neuronal activity ^51, 52, 53^.

Pvalb neurons also show cell-to-cell variability in the strength of the Kir-mediated inhibitory effect. Human Pvalb neurons with either high or low resting electrical resistance can show strong or weak Kir-mediated inhibition, but in the mouse, the cell with higher Rin showed a stronger Kir effect. This species difference may reflect the greater diversity of Pvalb neuron subtypes in the human compared to rodent neocortex ^41^. The strength of Kir activity may be specific to a subtype of neocortical Pvalb neuron, or the strength of Kir activity may be plastic and dynamically adjusted according to the activity history of the neurons ^47^.

### Kir channel types in human and mouse Pvalb interneurons

This study demonstrated a prominent Kir channel-mediated effect in human Pvalb neurons, similar to previous reports in rodent Pvalb neurons ^31^. Both species showed predominantly mRNA expression of Kir3.1 and Kir3.2 channels. These channel proteins were also detected in both human and mouse neurons by fluorescent immunohistochemical labeling at the extracellular membrane of soma. The results suggest that Kir3.1 and Kir3.2 channels are responsible for most of the Kir-mediated inhibition in the soma of Pvalb neurons in both species. The human neocortex displayed less diversity in the Kir channel types expressed in Pvalb neurons, which may have implications for therapeutic effect of Kir-channel modulators on these neurons ^33, 34^.

Mouse Pvalb neurons also exhibited robust mRNA expression of *KCNJ9* (Kir3.3 channels) both in our patch-sequenced neurons and in the Allen Institute database, whereas its expression was low in human Pvalb neuron types. However, constitutive deletion of the Kir3.3 subunit in the rodent brain has minor phenotypic consequences, and its role in the brain is not well understood ^54^. Kir2 family mRNAs (*KCNJ4* and *KCNJ12*) were prominent in mice but less strongly expressed in humans in both the Allen Institute and our datasets ^31^. Although mRNA transcript levels failed to associate with strength of the Kir effect in neurons, it can be speculated that the more diverse expression of Kir channel types in mouse cells (high expression of Kir2.2, Kir2.3, Kir3.3, and Kir6.2, which are weakly expressed in humans) is related to their greater need for Kir channel conductance density (and hence Kir channel protein) in their plasma membrane.

### Physiological mechanisms that drive the Kir-mediated inhibition of human Pvalb neurons

Human and mouse Pvalb neurons exhibit decreases in excitability attributable to Kir activation, effectively dampening excitatory potentials in somata by synaptic inputs. This inhibitory effect in Pvalb neurons is caused by both a negative shift of Vm further away from the AP threshold during hyperpolarization and shunting inhibition of excitatory potentials in the somatic region ^37^. Computational model simulations showed that Kir channels at the somatic plasma membrane effectively shunt (short-circuit) excitatory EPSPs in human and mouse Pvalb neurons.

NGFs in the cortex elicit combined GABA-A and GABA-B receptor-mediated postsynaptic responses in several types of postsynaptic neurons, including fast-spiking Pvalb neurons ^43^. Single APs from NGFs, as demonstrated in both humans and rodents ^12^, evoke inhibitory postsynaptic potentials that include a GABA-B receptor-mediated component mediated by Kir potassium channels ^12, 44, 45, 55^. Activation of this synaptic inhibitory connection might be related to the regulation of slow-wave sleep and the alternation of active and quiescent states in the neocortex ^56^.

Common receptor mechanisms associated with Kir channel activation in Pvalb neurons include opioid receptors. Opioid receptors in the brain are regulators of pain sensation and mood, and they are important targets of drugs of abuse ^57^. Mu- and delta-subtype receptors are expressed in Pvalb interneurons, in which they induce a Kir channel-mediated decrease in excitability that causes disinhibition in cortical circuits ^30, 58^ in both mice and primates ^30^.

## Methods

### Ethics statement

All procedures were approved by the University of Szeged Ethics Committee, the Regional Human Investigation Review Board (ref. 75/2014), and the Scientific and Research Ethics Committee (ETT TUKEB) BM/25042-1/2024 and were conducted in accordance with the tenets of the Declaration of Helsinki.

### Human brain slices

Slices were prepared from samples of the frontal, temporal, or other cortical regions removed during surgical treatment targeting deep brain structures. Detailed information on neocortical regions and patient demographics is presented in Supplementary Table 1. Anesthesia was induced with intravenous midazolam (0.03 mg/kg) and fentanyl (1–2 μg/kg) following a bolus intravenous injection of propofol (1–2 mg/kg). Additionally, patients received 0.5 mg/kg rocuronium to facilitate endotracheal intubation. During surgery, patients were ventilated with a 1:2 mixture of O_2_/N_2_O, and anesthesia was maintained with sevoflurane. Following surgery, the resected tissue blocks were immediately immersed in an ice-cold solution containing (in mM) 130 NaCl, 3.5 KCl, 1 NaH_2_PO_4_, 24 NaHCO_3_, 1 CaCl_2_, 3 MgSO_4_, and 10 D-(+)-glucose aerated with 95% O_2_/5% CO_2_ within a glass container on ice inside a thermally isolated box (within 20 min) and transported from the operating room to the electrophysiology laboratory with continuous 95% O_2_/5% CO_2_ aeration. Slices (350 μm thick) were prepared from the brain tissue block using a vibrating blade microtome (Microm HM 650 V, Thermo Fisher Scientific,

Waltham, MA, USA) and then incubated at 22°C–24°C for 1 h in slicing solution. The slicing solution was gradually replaced with recording solution (storage chamber circulating solution volume of 180 mL) using a pump (6–7 mL/min). The recording solution was identical to the slicing solution except that it additionally contained 3 mM CaCl_2_ and 1 mM MgSO_4_.

### Mouse brain slices

Transversal slices (350 μm) were prepared from the somatosensory and frontal cortices of 5– 12-week-old heterozygous B6.129P2-Pvalbtm1(cre)Arbr/J mice (stock 017320, B6 PVcre line, Jackson Laboratory, Bar Harbor, ME, USA) crossed with the Ai9 reporter line to express the tdTomato fluorophore in Pvalb neurons to assist in cell selection ^15^.

### Whole-cell recordings

Recordings were performed in a submerged chamber perfused with recording solution (8 mL/min) maintained at 36°C–37°C. The cells were patched using visual guidance with infrared differential interference contrast video microscopy and a water immersion ×20 objective with an additional ×2-4 zoom. An appropriate dichroic mirror and filter (Cy3) was used for mouse tissue tdTomato signal epifluorescence. All recordings were completed within 30 min from entering the whole-cell mode. Micropipettes (5–7 MΩ) were filled with an intracellular solution with the following composition (in mM): 126 K-gluconate, 8 NaCl, 4 ATP-Mg, 0.3 Na_2_-GTP, 10 HEPES, and 10 phosphocreatine (pH 7.0–7.2; 300 mOsm) supplemented with 0.3% (w/v) biocytin for subsequent staining with fluorophore-conjugated streptavidin. Recordings were performed using a Multiclamp 700B amplifier (Axon Instruments, Scottsdale, AZ, USA). The recorded signal was low-pass–filtered online at a cutoff frequency of 6–8 kHz (Bessel filter). The series resistance and pipette capacitance were compensated in the current–clamp mode. All parameters were measured from at least five reproduced traces. Data were acquired using Clampex software (Axon Instruments), digitized at 35–50 kHz, and analyzed offline using NeuroExpress (version 24.c.02, developed by Attila Szucs) ^15, 59^, pClamp (version 10.5, Axon Instruments), Spike2 (version 8.1, Cambridge Electronic Design, Milton, UK), OriginPro (version 9.5, OriginLab Corporation, Northampton, MA, USA), and SigmaPlot (version 14, Grafiti, Palo Alto, CA, USA).

### Ion channel blockers

ZD7288 and BaCl_2_ (Sigma-Aldrich, St. Louis, MO, USA) were diluted in physiological extracellular solution and applied by wash-in.

### Tissue fixation and visualization of biocytin-filled neurons

Cells filled with biocytin were visualized using either Alexa 488-conjugated streptavidin (1:1000, Jackson ImmunoResearch, West Grove, PA, USA) or Cy3-conjugated streptavidin (1:1000, Jackson ImmunoResearch). After recording, the slices were immediately fixed for at least 12 h in 4% paraformaldehyde and 15% picric acid in 0.1 M phosphate buffer (PB; pH 7.4) at 4°C and then stored at 4°C in 0.1 M PB containing 0.05% sodium azide as a preservative. All slices were embedded in 20% gelatin and further cut into 50–60-μm-thick sections in ice-cold PB using a vibratome (Microm HM 650 V). The sections were rinsed in 0.1 M PB (thrice for 10 min each), cryoprotected in 10%–20% sucrose solution in 0.1 M PB, flash-frozen in liquid nitrogen, and thawed in 0.1 M PB. They were then incubated in 0.1 M Tris-buffered saline (TBS; pH 7.4) containing fluorophore-conjugated streptavidin for 2–3 h at 22°C–24°C. After washing with 0.1 M PB (thrice for 10 min each), the sections were covered in Vectashield mounting medium (Vector Laboratories, Burlingame, CA, USA), placed under a coverslip, and examined under an epifluorescence microscope at ×20–60 magnification with the appropriate excitation light filters and dichroic mirrors (Leica DM 5000 B, Leica, Wetzlar, Germany) ^15^. Sections used for cell reconstruction were further incubated overnight in conjugated avidin–biotin horseradish peroxidase (1:300; Vector Labs) in TBS (pH 7.4) at 4°C. The enzymatic reaction was visualized by the glucose oxidase–diaminobenzidine–nickel method using 3,3′-diaminobenzidine tetrahydrochloride (0.05%) as the chromogen and 0.01% H_2_O_2_ as the oxidant. Sections were further treated with 1% OsO_4_ in 0.1 M PB. After several washes in distilled water, the sections were stained with 1% uranyl acetate, dehydrated in an ascending series of ethanol concentrations, infiltrated overnight with epoxy resin (Durcupan, Merck, Darmstadt, Germany), and embedded on glass slides. Light microscopic reconstructions were conducted using the Neurolucida system (MBF Bioscience, Williston, VT, USA) with a ×100 objective (Olympus BX51, Olympus UPlanFI, Olympus, Tokyo, Japan). Images were collapsed in the z-axis for illustration.

### Immunohistochemistry

Free-floating sections were washed three times in TBS containing 0.3% Triton-X (TBST) for 1 h at 20°C–24°C. They were then moved to blocking solution (20% horse serum in TBST). All sections were incubated in primary antibodies diluted in TBST for three nights at 4°C, followed by incubation with the appropriate fluorochrome-conjugated secondary antibody solution (2% blocking serum in TBST) overnight at 4°C. The sections were first washed in TBST (thrice for 15 min each), washed in 0.1 M PB (twice for 10 min each), and finally mounted on glass slides with Vectashield mounting medium. The following primary antibodies were used for immunostaining of human and mouse brain sections for confocal microscope studies: goat anti-pv (1:1000, Swant, Switzerland, www.swant.com), rabbit Anti-GIRK1 polyclonal (1:200, #APC-005, www.alomone.com), and rabbit Anti-GIRK2 antibody monoclonal (1:200, EPR23841-83, https://www.abcam.com/en-us/products/primary-antibodies/girk2-antibody-epr23841-83-ab259909#). Secondary antibodies: DARb Alexa 647–conjugated donkey anti-rabbit (1:200, Abcam, www.abcam.com), and DAGt Cy3-conjugated donkey anti-goat (1:400, Jackson ImmunoResearch www.jacksonimmuno.com). For dSTORM microscopy rabbit Anti-GIRK1 polyclonal was used (1:200, #APC-005, https://www.alomone.com/p/anti-kir3-2-girk2/APC-006).

### Immunofluorescence analysis

Brain slices labeled for immunofluorescence were acquired using a Leica Stellaris 8 laser-scanning confocal microscope (Nikon CFI Apo TIRF 100XC Oil, NA = 1.49) with 554- and 649-nm lasers for excitation of the fluorophores. The utilized emission filters for these lasers were 559–654 and 654–750 nm, respectively. Images were acquired using an HC PL APO CS2 ×63/1.40 oil immersion objective in the unidirectional scanning mode. Line analysis of the immunofluorescence intensity was then performed offline using LAS X Life Science Microscope Software (Leica) as described previously ^15^. For data pooling in Fig. 4, the mean absolute intensity values (ordinate) are presented. Confocal immunofluorescence microscopy images of the neurons were analyzed using ImageJ (US National Institutes of Health, Bethesda, MD, USA). To determine the edge and centroid of the cell soma, the images were segmented by adjusting the threshold of the Pvalb/tdTomato fluorescence signal. Segmentation was performed using the IsoData thresholding method. A straight line of the selected length was drawn at equal angular intervals. The intensity profiles of both Kir3.1 or Kir3.2 and

Pvalb/tdTomato fluorescence channels were acquired along each line (from outside to inside the cell). Segments extending from the distal end of each line to the centroid of the soma were aligned by the extracellular membrane location defined as 0-point by the Pvalb/tdTomato fluorescence signal threshold [(threshold = average of the cell background pixels fluorescence subtracted by the average of the cell pixels fluorescence) / 2] along the line when moving from extracellular to intracellular direction in line analysis (ImageJ). In the pooled analysis of lines, means and upper and lower quartiles were calculated for each position along the lines and normalized to the maximum average.

Super-resolution dSTORM measurements were performed on a custom-made inverted microscope based on a Nikon Eclipse Ti-E frame with an oil immersion objective (Nikon CFI Apo TIRF 100XC Oil, NA = 1.49), as described previously ^15^. Images of individual fluorescent dye molecules were captured by an iXon3 897 BV EMCCD camera (512 × 512 pixels with area size of 16×16 μm; Andor Technology, Belfast, Northern Ireland) with the following acquisition parameters: exposure time, 20 ms; EM gain, 200; and temperature, −75°C. Typically, between 20,000 and 25,000 frames were captured from a single region of interest. The Nikon Perfect Focus System was used to keep the sample in focus during the measurements. High-resolution images were reconstructed using rainSTORM localization software (http://laser.cheng.cam.ac.uk/wiki/index.php/Resources). Background autofluorescence was calculated as the weighted moving average of the neighboring pixel values and subtracted from each frame before the localization step. The pixel size of the final pixelized super-resolution images (typically 30 nm) was set according to the localization precision and localization density. The dSTORM experiments were conducted in GLOX switching buffer, and the sample was mounted onto a microscope slide. The imaging buffer was an aqueous solution diluted in PBS containing the GluOx enzymatic oxygen scavenging system, 2000 U/mL glucose oxidase (Sigma-Aldrich, catalog number: G2133-50KU), 40,000 U/mL catalase (Sigma-Aldrich, catalog number: C100), 25 mM potassium chloride (Sigma-Aldrich, catalog number: 204439), 22 mM tris(hydroxymethyl)aminomethane (Sigma-Aldrich, catalog number: T5941), 4 mM tris(2-carboxyethyl)phosphine (Sigma-Aldrich, catalog number: C4706) with 4% (w/v) glucose (Sigma-Aldrich, catalog number: 49139), and 100 mM β-mercaptoethylamine (Sigma-Aldrich, catalog number: M6500). The final pH was set to 7.4.

### Nucleus collection for patch-sequencing, cDNA amplification and library construction, and sequencing data processing

The protocols were described in detail elsewhere ^16^. To prepare the libraries, we used a protocol based on the smartseq2 procedure ^60, 61^. The study used 66-base paired-end reads that were aligned to GRCh38 using the annotation GTF file (GRCh38_r93) retrieved from Ensembl. Sequence alignment was performed using STAR v2.7.11a ^62^. The default parameters were used with the following arguments: --soloType SmartSeq --soloUMIdedup Exact -- soloStrand Unstranded --soloFeatures GeneFull. Full gene counts, including intron and exon counts, produced by STAR were used to calculate the TPM values, which were used for subsequent analyses.

### Identification of Pvalb neurons using single-nucleus transcriptomics analysis

The transcriptomic data of individual cells were uploaded to the Allen Institute Neocortical Neuron Type Identification System to categorize neocortical neuron types by the genes expressed [Allen Institute for Brain Science, 2024, MapMyCells, available from https://knowledge.brain-map.org/mapmycells/process/ (RRID:SCR_024672)], allowing Pvalb neuron identification in cells in which PVALB/Pvalb expression was below the detection limit. This algorithm uses a set of marker genes to predict the cluster/supertype of the cells based on their corresponding single-nucleus dataset. For human samples, the Seattle Alzheimer’s Disease Brain Cell Atlas reference taxonomy was used with the deep generative mapping algorithm, whereas mouse cells were classified using the 10X whole mouse brain reference taxonomy with the hierarchical mapping algorithm.

### Single-cell model

The program used in the simulations was written by Attila Szucs in the Delphi language ^59, 63^. The software uses Heun’s method of integration, and a fixed time step of 1 µs was used to integrate the differential equations. The soma model neuron was based on a previously described formalism ^42, 59^. The model featured a somatic compartment. G_leak_ of the somatic compartment in Fig.5 was 1.42 nS, G_Kir_ (maximum) was 10 nS (Fig. 6c1), and Cm of the somatic compartment was 80 pF. The leakage reversal potential was set to −68 mV. Parameters were chosen to replicate the basic response properties and carefully adjusted using the experimental data to attain the best match between the simulated and recorded voltage traces. All intrinsic voltage-dependent currents were calculated as previously described ^16^.

### Statistical analysis

The data were presented as the median and interquartile range. Statistical significance was tested using the Mann–Whitney U test, ANOVA on ranks (Kruskal–Wallis H test) with Dunn’s *post hoc* pairwise test, or one-way MANOVA with Bonferroni’s *post hoc* test to assess differences between multiple samples. Associativity and correlations were tested using the Spearman rank order method. Tests were performed using SigmaPlot 15. Inset symbols and icons were prepared from premade icons and templates using BioRender’s web-based free software (https://www.biorender.com).

**Supplementary Table 1.**
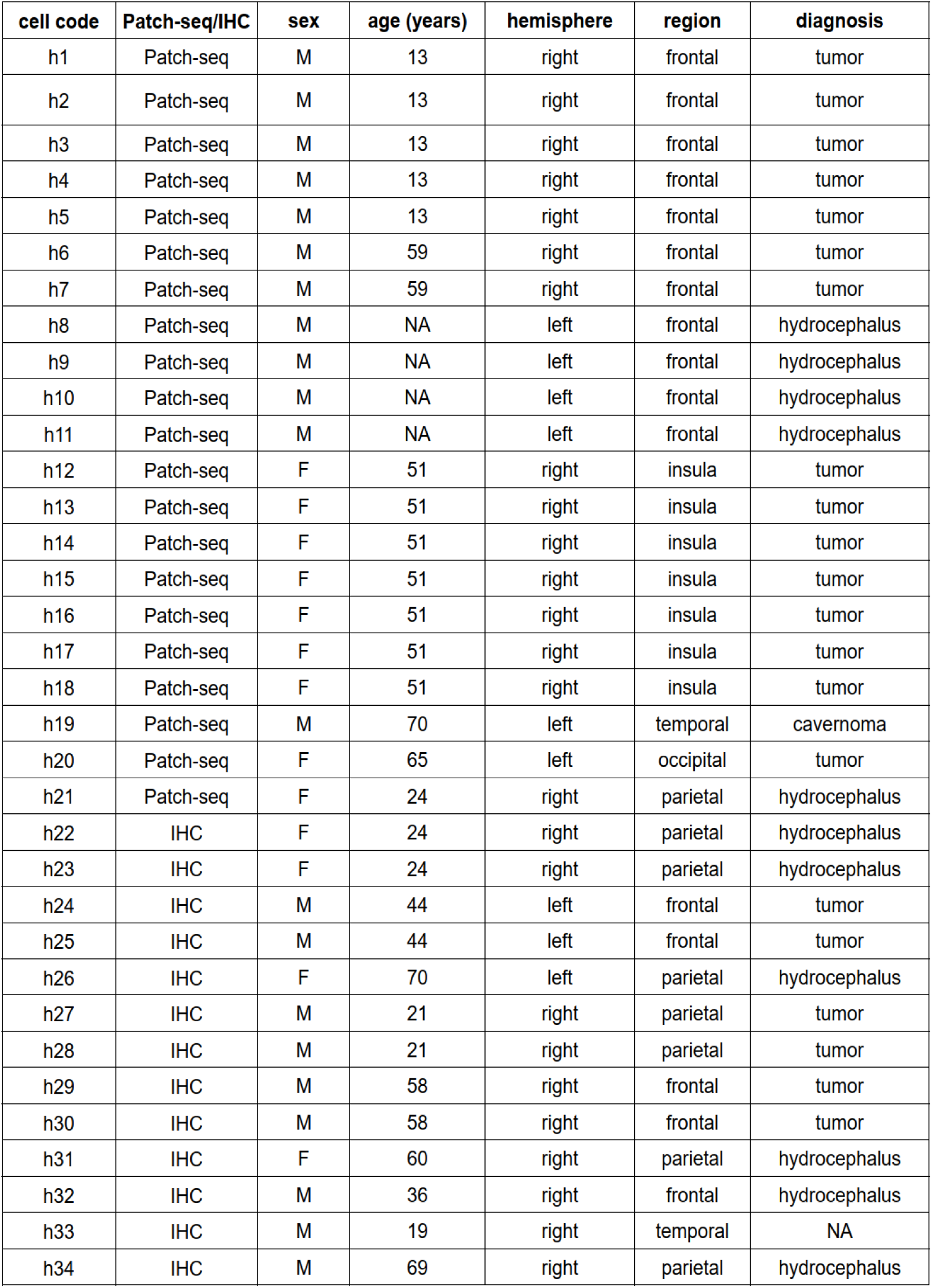
Details of human neocortical tissue resected during surgery, which was used to investigate the neurons in the study. From left to right: Cell code; Analysis method for parvalbumin (Pvalb) expression in the cells (IHC = immunohistochemistry; patch-seq = mRNA analysis); Patient sex (M = male, F = female) and age (years); Cerebral hemisphere (left or right); Cortical region; and Diagnosis for surgery.

**Supplementary Table 2.** A table presenting the bootstrapped p-values for a specific single-cell mRNA sequencing database containing individual cells obtained by patch-sequencing. These cells were categorized according to the cell type classification system from the Allen database (https://knowledge.brain-map.org/mapmycells/process/) [39]. The values in the table were generated by the MapMyCells algorithm, which compares the entire transcriptomic profile of our cells with that of the published database. In addition, the table contains the detected genes for each cell, expressed in thousands (k).

| HUMAN |  |  |  | MOUSE |  |  |  |
| --- | --- | --- | --- | --- | --- | --- | --- |
| cell ID | detected genes K | cluster label | bootstrapping p-value | cell ID | detected genes K | cluster label | bootstrapping p-value |
| h1 | 7.4 | Pvalb_2 | 1 | m1 | 3.2 | 114 | 1 |
| h2 | 5.1 | Pvalb_2 | 1 | m2 | 3.7 | 114 | 1 |
| h3 | 5.4 | Pvalb_2 | 1 | m3 | 3.5 | 114 | 1 |
| h4 | 7.1 | Pvalb_2 | 1 | m4 | 4 | 114 | 1 |
| h5 | 7.2 | Pvalb_2 | 0.76 | m5 | 6.8 | 114 | 1 |
| h6 | 4.4 | Pvalb_2 | 1 | m6 | 5.5 | 114 | 1 |
| h7 | 8 | Pvalb_2 | 1 | m7 | 4.6 | 114 | 0.78 |
| h8 | 3.8 | Pvalb_2 | 1 | m8 | 4.5 | 114 | 1 |
| h9 | 3.9 | Pvalb_2 | 1 | m9 | 4 | 115 | 1 |
| h10 | 6.6 | Chandelier_1 | 1 | m10 | 3.4 | 115 | 0.77 |
| h11 | 4.6 | Chandelier_1 | 1 |  |  |  |  |
| h12 | 8.3 | Pvalb_15 | 1 |  |  |  |  |
| h13 | 6.4 | Pvalb_15 | 1 |  |  |  |  |
| h14 | 8.3 | Pvalb_15 | 0.92 |  |  |  |  |
| h15 | 4.9 | Pvalb_15 | 1 |  |  |  |  |
| h16 | 7.9 | Pvalb_15 | 1 |  |  |  |  |
| h17 | 8 | Pvalb_15 | 1 |  |  |  |  |
| h18 | 4.2 | Pvalb_3 | 1 |  |  |  |  |
| h19 | 6.9 | Pvalb_8 | 0.96 |  |  |  |  |
| h20 | 6.8 | Pvalb_5 | 1 |  |  |  |  |
| h21 | 7 | Pvalb_5 | 1 |  |  |  |  |

### Data availability statement

All data needed to evaluate the conclusions in the paper are present in the paper and/or the Supplementary Materials. All datasets and custom codes used in this study are stored in Hungarian Center of Excellence for Molecular Medicine, Szeged, Hungary, and available on request from the corresponding author.

## Acknowledgements

K.L. declares support for the research of this work from the Ministry of Culture and Innovation of Hungary from the National Research (Project no. TKP-2021-EGA-05) provided by, Development and Innovation Fund financed under the TKP2021-EGA funding scheme, the Ministry of Culture and Innovation of Hungary from the National Research, Development and Innovation Fund, financed under the 2022-2.1.1-NL funding scheme (Project no. 2022-2.1.1-NL-2022-00005), EU’s Horizon 2020 research and innovation program under grant (No. 739593). Nemzeti Kutatási, Fejlesztési és Innovaciós Alap, Magyar Tudományos Akadémia - the National Brain Research Program Hungary (OTKA K 134279).

G.T. discloses support for the research of this work from The Hungarian Scientific Research Foundation (ANN-135291), Eötvös Loránd Research Network grants ELKH-SZTE Agykérgi Neuronhálózatok Kutatócsoport (KÖ-36/2021), Ministry of Human Capacities Hungary (20391-3/2018/FEKUSTRAT and NKP 16-3-VIII-3), National Research, Development and Innovation Office (GINOP 2.3.2-15-2016-00018), Élvonal (KKP 133807, ÚNKP-20-5 - SZTE-681 (G.T.), National Institutes of Health (U01MH114812 and UM1MH130981).

A.S. declares support for the research of this work from National Research, Development and Innovation Office (ANN_135291). The authors would like to thank Enago (www.enago.com) for the English language review.

## Author contributions

Conceptualization: K.L. and V.S. Methodology: A.D., A.T., A.S., B.B., B.K, D.W., G.H, G.T, J.L., M.E., P.B., and V.B. Investigation: A.D., A.S., A.T, D.W., E.B., G.M., J.L, S.F. and V.S. Data curation: A.D., A.T., E.S., and V.S. Formal analysis: A.D., A.T., B.K., E.B., S.F., and V.S. Supervision: K.L. Project administration: K.L. and V.S. Funding acquisition: K.L., G.T, A.S. Writing original draft: K.L. Writing review and editing: K.L.

